# Discovering functional sequences with RELICS, an analysis method for tiling CRISPR screens

**DOI:** 10.1101/687293

**Authors:** Patrick C. Fiaux, Hsiuyi V. Chen, Aaron R. Chen, Poshen B. Chen, Graham McVicker

## Abstract

CRISPR screens are a powerful new technology for the identification of genome sequences that affect cellular phenotypes such as gene expression, survival, and proliferation. By tiling single-guide RNA (sgRNA) target sites across large genomic regions, CRISPR screens have the potential to systematically discovery novel functional sequences, however, a lack of purpose-built analysis tools limits the effectiveness of this approach. Here we describe RELICS, a Bayesian hierarchical model for the discovery of functional sequences from tiling CRISPR screens. RELICS considers the overlapping effects of multiple nearby functional sequences, accounts for the ‘area of effect’ surrounding sgRNA target sites, models overdispersion in sgRNA counts, combines information across multiple pools, and estimates the number of functional sequences supported by the data. In simulations, RELICS outperforms existing methods and provides higher resolution predictions. We apply RELICS to published CRISPR interference and CRISPR activation screens and predict novel regulatory sequences, several of which we experimentally validate. In summary, RELICS is a powerful new analysis method for tiling CRISPR screens that enables the discovery of functional sequences with unprecedented resolution and accuracy.

## Background

CRISPR screens are an important class of new methods that perturb the genome to discover sequences that affect cellular phenotypes such as growth, survival, or gene expression. In a CRISPR screen, thousands of single-guide RNAs (sgRNAs) are delivered to cells and target sequences for mutation, repression, or activation. The cells are subsequently sorted into pools based on gene expression or subjected to selection. To target sequences for mutation, sgRNAs are introduced to cells alongside Cas9. Cas9 creates double strand breaks at the targeted sites, and mutations are introduced by error-prone non-homologous end-joining (1). In a CRISPR interference (CRISPRi) experiment, targeted sites are silenced by a deactivated Cas9 (dCas9) enzyme fused to a repressive domain such as the Krüppel-associated box (dCas9:KRAB) (2, 3). Similarly, in a CRISPR activation (CRISPRa) experiment, dCas9 is fused to an activation domain such as VP64 or p300 (4–6).

The first CRISPR screens targeted protein-coding genes (7), but the decreasing cost of oligo synthesis has recently enabled screens that tile sgRNAs across non-coding regions of the genome (8–18). These tiling CRISPR screens are particularly useful for the discovery of novel functional sequences, because they are not biased to sequences where function is detectable by existing assays such as those that measure histone modifications (e.g. H3K4me1 and H3K27ac) (19–21), open chromatin (22, 23), or enhancer RNAs (eRNAs) (24, 25). In addition, unlike massively parallel reporter assays (MPRAs) (26–28), tiling CRISPR screens measure the effects of targeted sequences in their native genomic context, and can connect discovered regulatory sequences to the genes that they control.

Tiling CRISPR screens have the potential to revolutionize our understanding of genome function, but their analysis presents several challenges. One challenge is that CRISPR effector proteins have an ‘area of effect’ that extends well beyond their target sites. In Cas9-based screens, targeting with a single sgRNA typically results in small indels within 20 bases of the target site (29), however, in CRISPRa or CRISPRi screens, the area of effect can extend over 1kb or more (3). The advantage of a large area of effect is that larger regions of the genome can be interrogated by increasing the spacing between sgRNA target sites, but the disadvantage is that it is more difficult to narrow down the causal genomic region.

Additional challenges in the analysis of tiling CRISPR screens include the inherent variability in sgRNA efficiencies (30–33), the noisy and overdispersed nature of genomic count data (34, 35), the need to combine information across multiple sgRNAs, and the possibility that sgRNAs affect multiple functional sequences near their target sites. Furthermore, CRISPR screens often generate counts from multiple pools (e.g. pre-/post-selection or expression pools), however, existing analysis methods are not capable of analyzing all of the pools at once and instead compare two pools at a time.

Here we describe RELICS (**<u>R</u>**egulatory **<u>E</u>**lement **<u>L</u>**ocation **<u>I</u>**dentification in **<u>C</u>**RISPR **<u>S</u>**creens), a new method for the analysis of CRISPR screens which uses a flexible Bayesian hierarchical model to jointly analyze sgRNA counts across multiple pools. RELICS considers the overlapping effects of multiple sgRNAs and estimates the number of functional sequences that are supported by the data. We apply RELICS to a large number of simulated datasets and find that it is the best-performing analysis method under a wide variety of conditions. In addition, the sequences predicted by RELICS have higher resolution—they are smaller but still contain the true functional sequences. We confirm this improved accuracy by running RELICS on published CRISPRa and CRISPRi datasets and experimentally validating the predictions.

## Results

### The RELICS model

RELICS is a flexible Bayesian hierarchical model for the discovery of functional sequences from pooled CRISPR screens including those based on cell dropout, cell proliferation and gene expression (Fig. 1a,b). The goals of RELICS are to determine how many functional sequences are supported by the data, and to determine the location of these functional sequences in the genome. To accomplish these goals, RELICS divides the screened sequence into small genome segments and associates each sgRNA with the genome segments that overlap its area of effect (Fig. 1c). RELICS assumes that genome segments containing functional sequences affect the counts of the overlapping sgRNAs. For example, an sgRNA that targets a gene’s promoter might be expected to have a higher proportion of its counts in a low expression pool compared to sgRNAs that target non-functional sequences. RELICS learns how functional sequences affect sgRNA counts and then uses the observed sgRNA counts to calculate the posterior probability that a genome segment, *m*, contains a functional sequence, *k*. We refer to the probability that a genome segment contains a functional sequence as the ‘functional sequence probability’ and denote it *π_k,m_*.

**Figure 1.**
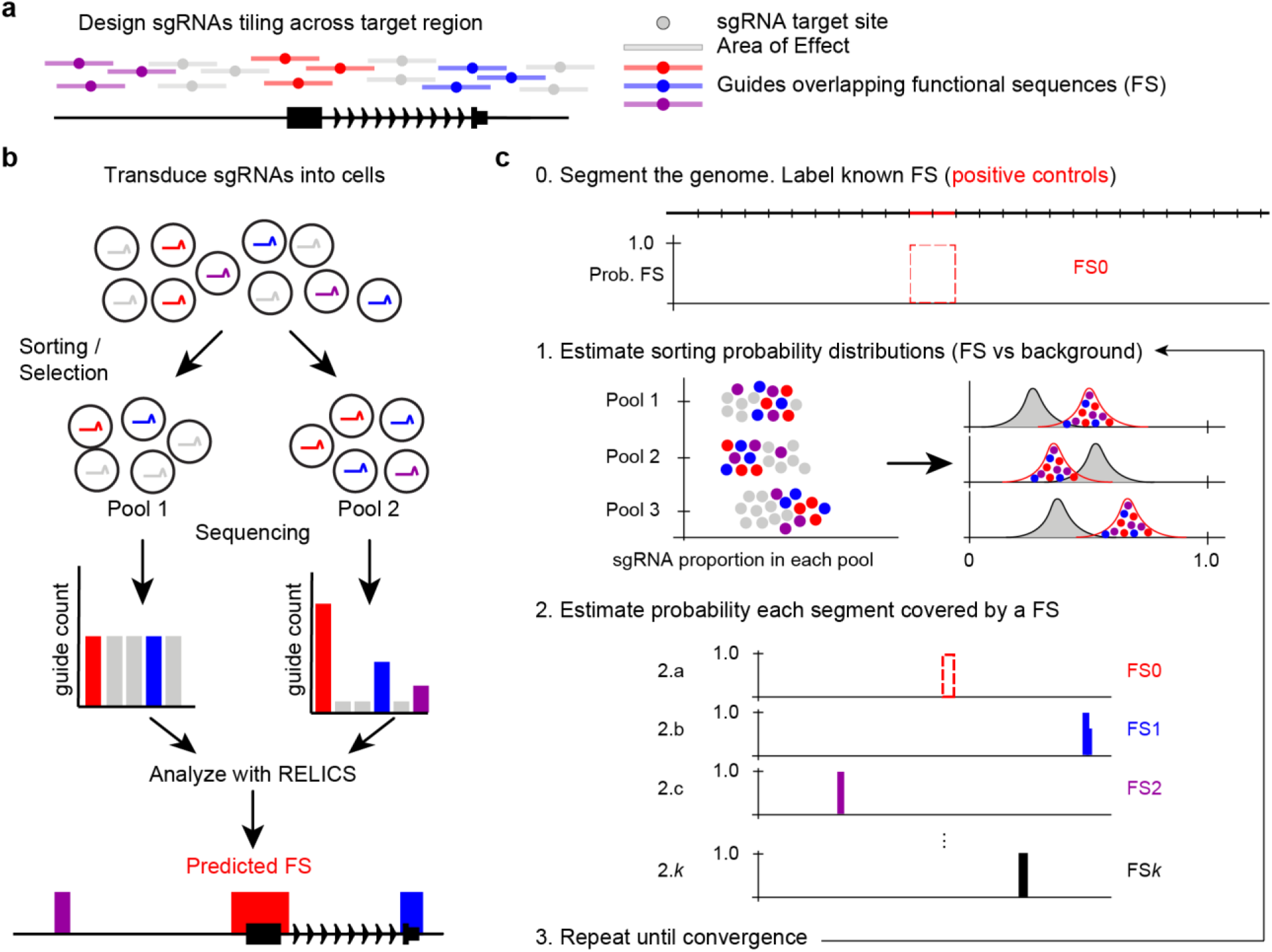
Schematic of a tiling CRISPR screen and RELICS. **(a)** sgRNAs are designed to target sites tiling across a genomic region. Bars around each target site indicate the area of effect. sgRNAs ‘overlap’ genome sequences within their area of effect. In this example, sgRNAs overlapping functional sequences (FSs) are colored red, purple and blue. **(b)** In a CRISPR screen, sgRNAs are introduced into cells and the cells are sorted into pools or subjected to selection (e.g. for proliferation or survival). The sgRNAs in each pool are counted and analyzed by RELICS to predict FSs. **(c)** RELICS predicts FSs using an Iterative Bayesian Stepwise Selection Algorithm. <u>Step 0 (initialization):</u> the screened region is divided into small (e.g. 100bp) genome segments; segments containing known FSs are labeled FS0. <u>Step 1:</u> Using the predicted FSs, RELICS estimates sorting probability distributions for sgRNAs that do/do-not overlap FSs. <u>Step 2:</u> RELICS uses the sorting probability distributions to compute the probability each genome segment contains a functional sequence. <u>Step 3:</u> Steps 1-3 are repeated until convergence.

RELICS models the observed sgRNA counts across pools using a Dirichlet multinomial distribution, which allows multiple pools to be analyzed while controlling for extra variability (overdispersion) in the data. The Dirichlet portion of the distribution describes the probability that a cell containing an sgRNA will be sorted into each pool. RELICS uses different Dirichlet distributions to describe the sorting probabilities for sgRNAs that do or do not overlap functional sequences (Fig. 1c). RELICS estimates the hyperparameters for both Dirichlet distributions empirically by maximum likelihood. This process is assisted by specifying known functional sequences (positive controls), which we designate functional sequence 0 (FS0). For example, in a CRISPRa or CRISPRi gene expression screen, genome segments overlapping the promoter of the target gene can be labeled as FS0. Note that the hyperparameters are estimated not only from FS0, but also from the placement of all other functional sequences (FS1 through FSK).

Using this model, RELICS can calculate the likelihood of any configuration (positions and sizes) of functional sequences on the genome sequence. These likelihoods can be used (in combination with prior probabilities), to calculate the functional sequence probability, *π_k,m_*, for each genome segment. Exact calculation of *π_k,m_* is impractical, however, because likelihoods must be calculated for every possible functional sequence configuration. To overcome this problem, RELICS uses a novel Iterative Bayesian Stepwise Selection (IBSS) algorithm (36) to calculate approximate functional sequence probabilities. RELICS’ IBSS algorithm assumes there are *K* functional sequences (of variable length), and places them one at a time, while taking into account the placements of all of the other functional sequences. After placing the final functional sequence, the entire process is repeated until convergence. This algorithm is efficient and allows for uncertainty in both the location and size of each functional sequence.

A key parameter in RELICS is the number of functional sequences, *K*, which must be specified prior to running the IBSS algorithm. Since the true number of functional sequences is unknown, we developed a procedure to automatically estimate *K* from the data, based on the observation that the predicted functional sequences begin to overlap when *K* is too high (see Methods). We use this automated estimation of *K* in all applications of RELICS.

The final output from RELICS is an estimate of *K*, the number of functional sequences that can be identified from the data, and a matrix ***π***, which gives the probability that each genome segment contains each functional sequence. An important feature of RELICS is that it outputs separate probabilities for each functional sequence, providing discrete genome segments that can be used for follow-up validation experiments (Fig. 1c). We plot these functional sequence probabilities as separate tracks (Figs. 1c, 2a, 4a) or as a combined track with different colors (Figs. 4b, 4c, 5a). In addition, the functional sequences output by RELICS are rank-ordered, with functional sequence 1 (FS1) having the strongest statistical support.

**Figure 2.**
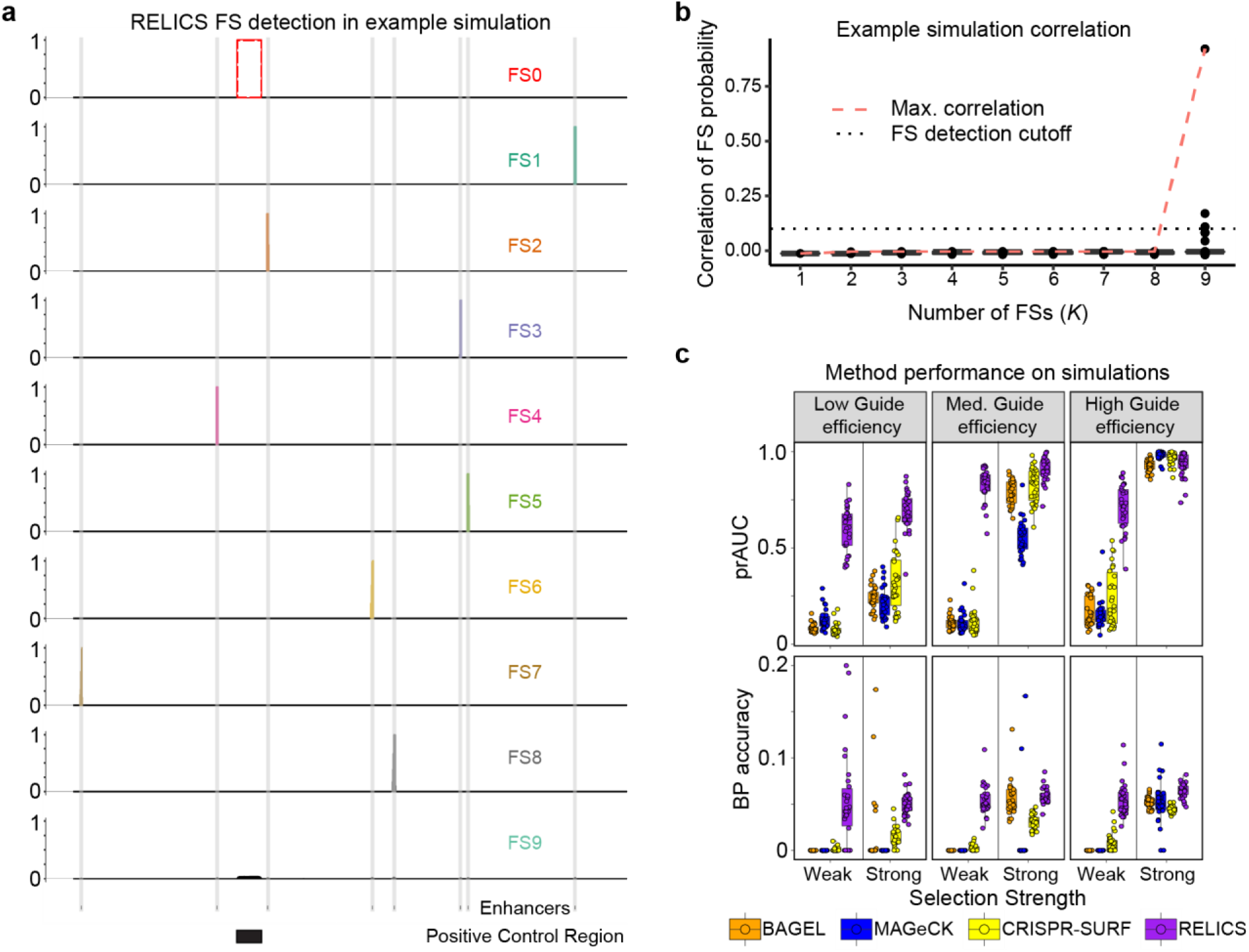
RELICS functional sequence (FS) prediction and performance of RELICS and other analysis methods on simulated data. **(a)** Example of functional sequence (FS) probabilities output by RELICS for a single simulation. FS0 is the known positive control sequence. RELICS correctly predicts all 8 of the simulated enhancers (grey vertical bars), which it labels FS1-FS8. FS9 is displayed to demonstrate the choice of *K* = 8. **(b)** Boxplots of pairwise correlations of FS probabilities for different choices of *K* (number of FSs). The red dashed line is the maximum pairwise correlation. The black dotted line is the cutoff used to determine *K*. In this example, RELICS chooses *K* = 8. The hinges of the boxplots correspond to the first and third quartiles, the center lines are the medians, and the whiskers extend to the furthest datapoints that are within 1.5x the interquartile range from the hinge. **(c)** Performance on 180 simulations summarized with area under the precision-recall curve (prAUC) and base pair (BP) accuracy. For this figure, 30 simulations were performed for 6 simulation scenarios with high read depth and different values of the key parameters “selection strength” and “guide efficiency”. Additional simulations and performance metrics are provided in Supplementary Figures 2 and 3. BP accuracy is the fraction of the significant bases identified by each method which are true functional sequences. In these simulations, all methods predict regions that are substantially larger than the small (50bp) simulated functional sequences due to the comparatively large area of effect around sgRNA target sites (1000bp), RELICS predicts the smallest regions with the highest BP accuracy.

**Figure 3.**
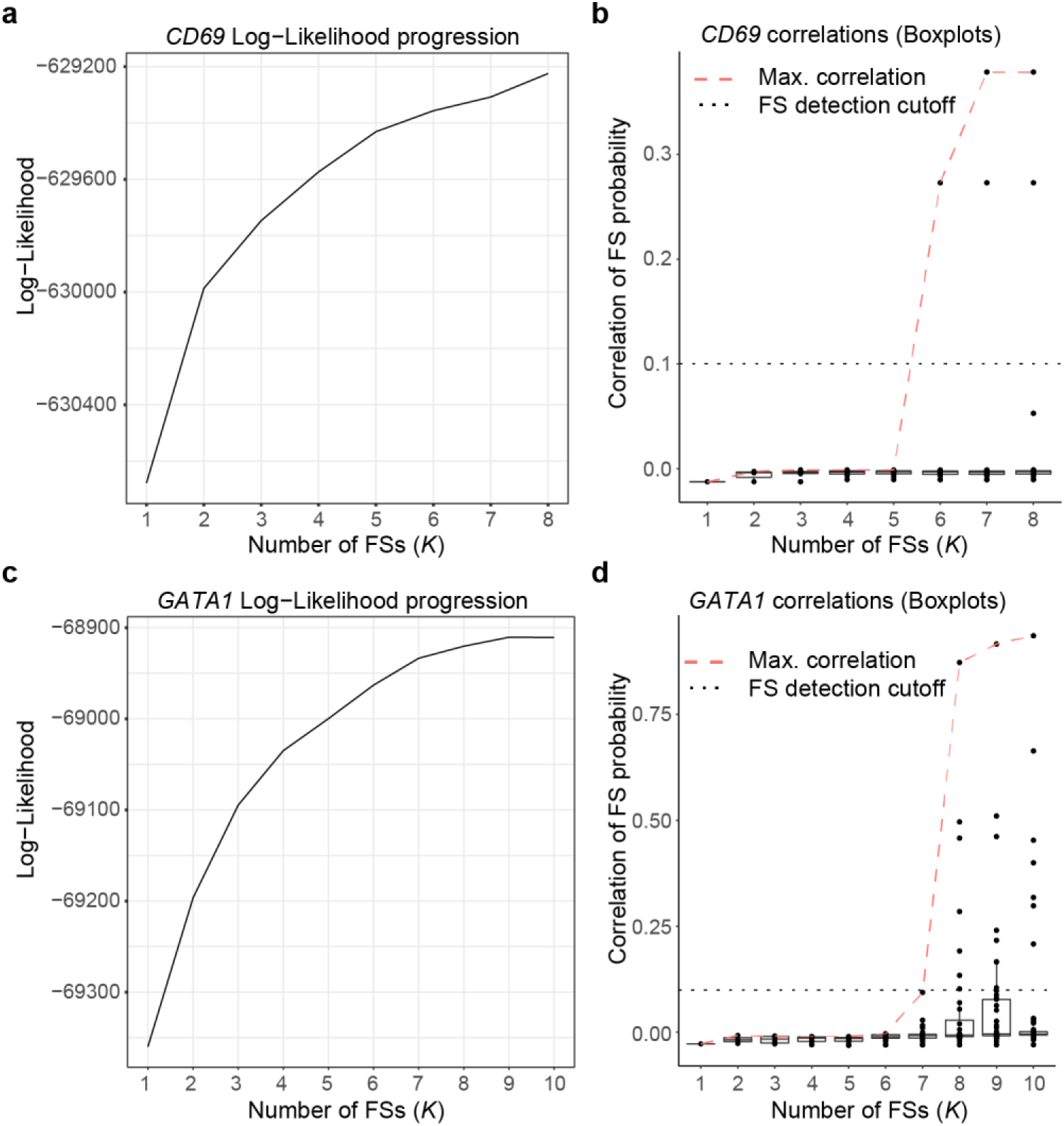
Log-likelihood progression and pairwise correlations between functional sequence (FS) probabilities as the number of functional sequences (*K*) is increased. **(a)** Log likelihood as the number of FSs increases for *CD69*. **(b)** Pairwise FS placement probability correlations for *CD69*. The red dashed lines indicate the highest pairwise correlation across all FSs. **(c)** Log-likelihood progression for *GATA1*. (d) Pairwise FS placement probability correlations for *GATA1*. The hinges of the boxplots in (b) and (d) correspond to the first and third quartiles, the center lines are the medians, and the whiskers extend to the furthest datapoints that are within 1.5x the interquartile range from the hinge.

**Figure 4.**
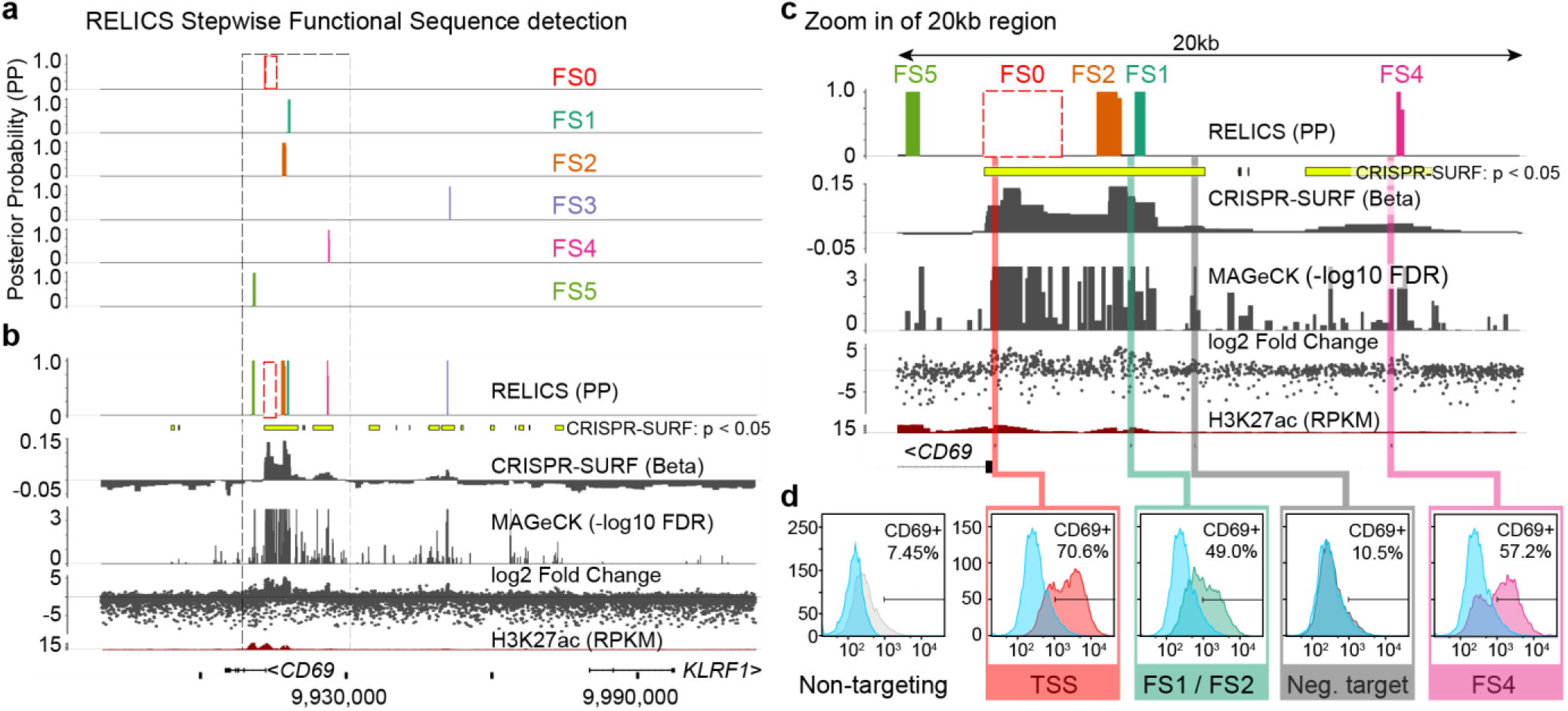
Analysis of a published CRISPR activation (CRISPRa) screen for *CD69* expression in Jurkat T cells. **(a)** RELICS detects 5 functional sequences (FSs) labeled FS1-FS5. FS0 is a known positive control sequence (the CD69 promoter) provided as input to RELICS and CRISPR-SURF. **(b)** Analysis of the CD69 screen by RELICS, CRISPR-SURF, MAGeCK and log2 fold change. The RELICS probabilities for each FS are collapsed into a single track. An H3K27ac ChIP-seq track for Jurkat cells is included. **(c)** Zoom in of a 20kb region (indicated by dashed box in (a) and (b)). Experimentally tested regions are indicated by colored bars. **(d)** Experimental validations. Lentiviruses carrying sgRNAs targeting different regions were transduced to Jurkat cells expressing dCas9:VP64, and the expression of CD69 was measured by flow cytometry. The results from each experiment are overlaid those from a non-targeting negative control sgRNA (blue). Targeting sgRNAs were chosen for their high predicted efficiency and in some cases are adjacent to the predicted FS rather than within the FS. Results from additional validation experiments are shown in Supplementary Figure 4.

**Figure 5.**
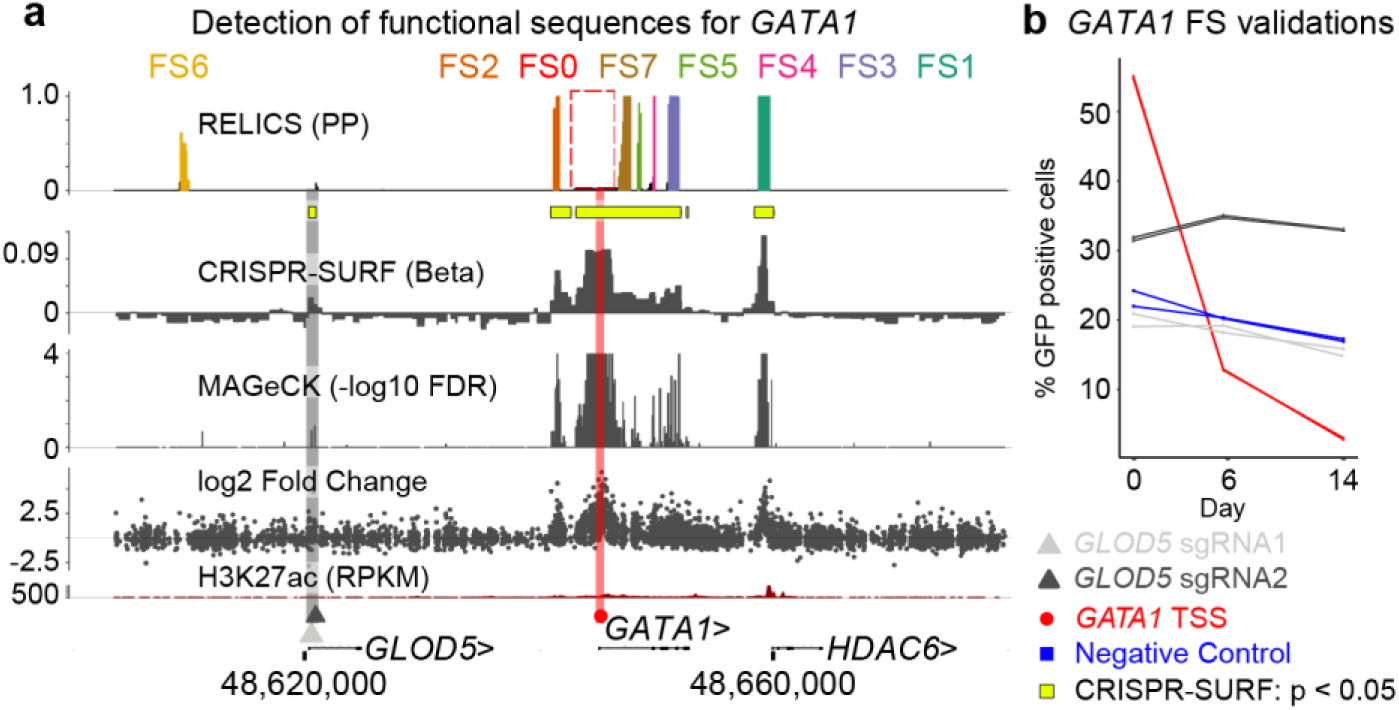
Analysis of a published CRISPR inhibition (CRISPRi) cellular proliferation screen around *GATA1*. **(a)** Results from RELICS and other analysis methods. RELICS detects 7 functional sequences (FS1-7). FS1 and FS2 have previously been validated; FS7, FS5, FS4 and FS3 fall within GATA1. sgRNA target sites for validation experiments are indicated with a red circle (GATA1 promoter) and grey and black triangles (*GLOD5*). **(b)** Results from validation experiments using sgRNAs targeting sites indicated in panel A. Each validation experiment is a cellular proliferation assay, in which the percent of GFP-positive cells (i.e. those that received the sgRNA) are measured at day 0, day 6 and day 14. While targeting the *GATA1* promoter greatly reduces proliferation, targeting the *GLOD5* region does not change proliferation compared to a negative control sgRNA, which targets a non-functional region on another chromosome.

### Application of RELICS to simulated data

To evaluate the performance of RELICS and other analysis methods, we developed a CRISPR screen simulation framework, CRSsim, that generates realistic datasets where the ground truth is known (Supplementary Figure S1). The simulated datasets are useful to benchmark performance since there is currently no large gold-standard set of known functional sequences. We simulated datasets with two different sequencing depths (medium/high), three sgRNA efficiency distributions (low/medium/high), and two selection strengths (weak/strong). The selection strength describes the probability of a cell being sorted into each pool given the presence or absence of an sgRNA targeting a functional sequence. We performed 30 simulations for the 12 combinations of parameter settings under two experimental scenarios: scenario 1 a cellular proliferation screen with two pools per replicate, and scenario 2 a gene expression screen with four pools per replicate. In total we performed 720 simulations, with each simulation consisting of 8,700 sgRNAs targeting a 150kb region with 8 functional sequences each spanning 50bp.

We ran RELICS on the simulated datasets and allowed it to estimate the number of functional sequences, *K*, by calculating the pairwise correlation between functional sequences probabilities for different values of *K*. An example of RELICS’ predictions on a single simulated dataset is shown in Figure 2. In this example, RELICS identified all 8 of the functional sequences before the pairwise correlations between functional sequence probabilities increased due to overlapping placements (Fig 2a, b).

We compared RELICS’ predictions on simulated data to those from (i) log fold change; (ii) MAGeCK(37), the dominant method for gene-based CRISPR screens; (iii) BAGEL (38), a supervised method for gene-based CRISPR screens; and (iv) CRISPR-SURF (39), a new tool for tiling CRISPR screens. We first quantified performance using scores output by each method and computing the area under the precision-recall curve (prAUC). As scores, we used the functional sequence probability (RELICS), the effect size (CRISPR-SURF), the BayesFactor (BAGEL), and the −log10(FDR) (MAGeCK). We found that RELICS has by far the best performance with the highest prAUC in 306/360 simulations for scenario 1 and 229/360 simulations for scenario 2 (Fig 2c, Supplementary Figures S2a, S3a). We next quantified performance using the functional sequences that each method reported as significant. RELICS had the highest precision in 308/360 of the simulations in scenario 1 and 291/360 of the simulations in scenario 2 (Supplementary figure S2b, S3b). RELICS also had the highest recall in 285/360 of the scenario 1 simulations and 357/360 of the scenario 2 simulations (Supplementary figure S2c, S3c). Notably, RELICS has far better performance than the other methods in scenarios with low sgRNA efficiency and/or weak selection strength. This is important because many sgRNAs in screening libraries have low efficiency (31).

We quantified the resolution of predicted functional sequences with basepair (BP) accuracy—the total number of significant genomic bases that overlap with true functional sequence. Due to the large area of effect (1000bp) and small functional sequences (50bp) used by the simulations, all of the methods had low BP accuracy and predicted regions that are substantially larger than the functional sequences. Nonetheless, the predictions made by RELICS covered much smaller regions than the other methods and it had the highest BP accuracy in 301/360 of the scenario 1 simulations and 327/360 of the scenario 2 simulations (Fig 2c, Supplementary figures S2d, S2d). Note that these comparisons are not biased to favor RELICS since the simulations were designed to mimic experimental procedures and are not tailored to RELICS. In summary, RELICS outperforms the other methods on simulated data and has the best overall precision, recall, and resolution.

### Application of RELICS to published datasets

We applied RELICS to published tiling CRISPRa screens for functional sequences that affect the expression of *CD69* and *IL2RA* in Jurkat T cells (15). For both genes, cells were flow sorted into 4 pools based on expression (negative, low, medium, high), and in the published analysis putative functional sequences were identified by computing log fold change between pairs of pools. We ran RELICS, CRISPR-SURF, and MAGeCK on both datasets. RELICS predicts *K* = 5 functional sequences for *CD69* (Fig. 3a, b), which we label as FS1 through FS5 (Fig. 4). In contrast, MAGeCK predicts a large number of significant regions (50) under a false discovery rate of 0.05, while CRISPR-SURF predicts multiple large functional sequences at and downstream of the *CD69* promoter (Fig. 4b). To test a subset of the predictions, we designed sgRNAs to target these predictions, cloned them into lentiviral vectors, and transduced them into Jurkat T cells expressing dCas9:VP64. As a negative control we used a non-targeting sgRNA, and as positive control we used an sgRNA targeting a site near the *CD69* transcription start site (Fig. 4c,d). The region including FS1 and FS2 was previously found to activate *CD69* expression (15) and we confirm that CRISPRa targeting of this region increased the number of *CD69* positive cells. Similarly, we validated that FS4 also increases *CD69* expression. All methods detected a signal at FS3, however we were not able to confirm that targeting of this region affects *CD69* expression (Supplementary Fig S4). Finally, targeting a “negative sequence” (NS) that was not predicted by RELICS, but was predicted by MAGeCK and is contained within a large significant region reported by CRISPR-SURF did not change *CD69* expression. These results confirm that the predictions made by RELICS are accurate, even when they are discordant with the other methods. In addition, the output from RELICS is easier to interpret because it provides ranked, cleanly-delineated predictions for each functional sequence (Fig. 4a).

RELICS identified 16 functional sequences for *IL2RA*, all of which are within or very close to the 6 regions identified in the original study (Supplementary Fig. S5). Since RELICS predictions have higher resolution than other methods (Fig. 2d, 4c, d, Supplementary Fig. 2d, 3d), the multiple smaller regions predicted by RELICS may reflect the true presence of multiple functional sequences. Alternatively, some large functional sequences may be split into smaller sequences by RELICS, particularly if some of the sgRNAs targeting the middle of the sequence have very low efficiency.

We next applied RELICS to a CRISPRi proliferation screen surrounding the *MYC* locus in K562 cells (9). As with *IL2RA*, RELICS detected all of the previously-reported signals, but split several of them into smaller predicted functional sequences. Interestingly, RELICS also identified 5 regions that have not been previously reported. One example is FS16, which is just 3kb upstream of the previously reported “e2” element. Other examples are FS7 and FS14, which both fall within an intron of the non-coding RNA *LINC00824* (Supplementary Fig. S6).

Finally, we applied RELICS to a CRISPRi proliferation screen surrounding *GATA1* (9). RELICS predicted 7 functional sequences (Fig. 3c, d), two of which (FS1 and FS2) have been previously validated (9), four of which are within *GATA1* (FS3, FS4, FS5, FS7), and one of which is upstream of *GLOD5* (FS6) (Fig. 5a). The original study predicted a functional region near *GLOD5* that CRISPR-SURF also considered to be significant (Fig. 5a). While RELICS found some signal at this region, it reported a low functional sequence probability (*π*<0.1 for all genome segments in this region). This suggested to us that the region near *GLOD5* may be a false positive detected by the other methods. To test this hypothesis, we used CRISPRi to target the *GLOD5* region, the *GATA1* promoter (positive control), and a sequence on chromosome 8 (negative control). While the sgRNAs targeting the *GATA1* promoter decreased cellular proliferation as expected, targeting of the *GLOD5* region did not affect proliferation relative to the negative control (Fig. 5b). Thus, this region is unlikely to be a functional sequence, and the low probability reported by RELICS is appropriate. Overall, RELICS accurately identifies functional sequences, and provides an easily interpretable output that quantifies its certainty that a region contains a functional sequence.

## Discussion

We have developed RELICS, a Bayesian hierarchical model for the analysis of CRISPR screens. Unlike gene-based analysis methods, RELICS is specifically designed to analyze tiling CRISPR screens where the locations of functional sequences are not known. The RELICS model provides numerous advantages. First, it considers the overlapping effects of multiple nearby sgRNA target sites and functional sequences. Second, it provides interpretable probabilistic output for each functional sequence that can be used to delineate small genome regions that confidently contain each functional sequence. Third, it models sgRNA counts appropriately without requiring transformation of the data or assuming normality. Fourth, it increases power by jointly modeling data from multiple sgRNA pools. While other methods include some of these features (e.g. CRISPR-SURF deconvolves the effects of multiple sgRNAs, and MAGeCK models overdispersed count data with a negative binomial distribution), only RELICS combines all of them into a single model. We also tested the robustness of RELICS to sequencing depth and density of sgRNA target sites by downsampling 75% of the sgRNA counts as well as 75% of all sgRNAs for the *CD69* CRISPRa screen (Supplementary Fig. S7). In both cases, the RELICS predictions remained consistent indicating that it is relatively insensitive to sequencing depth and sgRNA density. This is also supported by our simulations, which found that sgRNA efficiency and functional sequence strength affect performance more than sequencing depth (Supplementary Figs S2, S3).

To test the performance of RELICS and other analysis methods we developed a simulation tool, CRSSim. Using CRSsim, we generated hundreds of datasets, performed the first systematic comparison of analysis methods for tiling non-coding CRISPR screens, and found that RELICS has the best performance under a wide variety of conditions. CRSsim is open-source and we envision that it will be a useful tool for future performance comparisons, power analyses, and making informed decisions about experimental designs for CRISPR screens (e.g. spacing of sites targeted by sgRNAs, the sequencing depth, and the complexity of the vector library).

RELICS leverages known functional sequences (labeled positive controls) as well as unlabeled sequences to learn model hyperparameters. It is therefore a semi-supervised method. CRISPR-SURF and BAGEL similarly make use of known control sequences. A strength of this approach is that RELICS learns the behavior of sgRNAs across different pools from the data. A limitation is that it might be difficult to apply RELICS to datasets that do not contain a known positive control region or that have a low number of sgRNAs overlapping positive control regions. In addition, positive controls may not adequately represent all types of functional sequences, such as repressive or weak regulatory elements. As an alternative approach, the hyperparameters can be specified by the user. It may also be possible to develop an unsupervised learning approach where positive control labels are not provided and instead sequences with similarly-behaving sgRNAs are identified by clustering. Ideally the categories identified by such an approach would represent different types of sequences (e.g. strong regulatory elements, weak regulatory elements, silencers, non-regulatory elements).

RELICS does not currently model variation in sgRNA efficiency, variation in the strength of regulatory elements, or off-target effects. These limitations are mitigated by combining information across multiple sgRNAs, and by modeling overdispersion, which allows for the extra-variability that is introduced by factors such as off-target effects (31, 40) or sgRNA efficiency (30, 32, 33). Our simulations demonstrate that RELICS performs better compared to other methods, especially in the presence of weak enhancers or weak sgRNA efficiency (Fig. 2c, Supplementary Figs. S2, S3). Nonetheless, future versions of RELICS that explicitly model these sources of variation will likely achieve even better results.

In summary, RELICS performs better than previous analysis methods under most simulation conditions and, when applied to real data, identifies both previously-validated functional sequences and new functional sequences. Thus, RELICS is an extremely useful tool for the discovery of functional sequences from tiling CRISPR screens.

## Methods

### Simulation framework

The CRSsim simulation tool is open source and available on GitHub (https://github.com/patfiaux/CRSsim). To run it, the user first specifies the screen type, which can either be a selection screen (e.g. drop out or proliferation) or an expression screen, with an arbitrary number of expression pools. Next, the user specifies the screening method (Cas9, CRISPRi, CRISPRa, dual-guide CRISPR), the number and size of true functional sequences, the spacing/density of sgRNA target sites, and the distribution of sgRNA counts in the initial set of cells (i.e. the frequency of each sgRNA).

The probability of a cell being sorted into each pool depends on whether the sgRNA in that cell targets a functional sequence. We refer to the difference in sorting probabilities between sgRNAs that do/do not target functional sequences as the “strength of selection”, *T*. The sorting of sgRNAs into the different pools is simulated by sampling from a Dirichlet multinomial distribution. The sorting probabilities of sgRNAs that do not target functional sequences are given by a vector of Dirichlet parameters *α*, where the probability of being sorted into pool *j* is 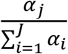. The sorting probabilities of an sgRNA, *n*, that overlaps a functional sequence, *k*, depends on the strength of selection, *T*, the sgRNA efficiency, *f_n_*, and the strength of the functional sequence, *h_k_*. Both *f_n_* and *h_k_* are proportions between 0 and 1. Guides targeting strong functional sequences, with a strength near *h_k_* = 1, behave like positive control sgRNAs. Guides targeting non-functional sequences have *h_k_* = 0. The simulation sets the efficiency of each sgRNA and the strength of each functional sequence by sampling from beta distributions with user-configurable shapes. Finally, the user can specify how deeply the sgRNA pools are sequenced.

The sgRNA efficiency, *f_n_*, specifies the fraction of cells where the sgRNA is ‘effective’ and perturbs the sorting probabilities. For example, if *c* cells contain sgRNA *n*, then *w* = *cf_n_* cells contain sgRNAs that are effective and they are sorted with probabilities specified by the Dirichlet vector *α* + *h_k_T*. The sorting vector for the c–*w* cells with ‘ineffective’ sgRNAs, is simply *α*.

Each simulation starts with an sgRNA library with a distribution of sgRNA counts. We assume the sgRNA counts follow a zero-inflated negative binomial distribution (ZINB), *y*~*ZINB*(*μ, d, ε*), where *μ* is the mean, *d* is the dispersion and *ε* is the proportion of the distribution that comes from the zero point mass. The parameters of the ZINB distribution can be specified or estimated by maximum likelihood from a provided table of sgRNA counts. After the sgRNA counts in the sgRNA library are obtained by sampling from the ZINB distribution, an input pool of cells containing sgRNAs is generated by performing multinomial sampling from the sgRNA library. We simulate the sorting of cells from the input pool into downstream pools by sampling from the Dirichlet multinomial distribution described above. Lastly, we simulate the sequencing step and obtain the count of sgRNAs in the sorted pools by drawing from a multivariate hypergeometric distribution (sampling without replacement) or a multinomial distribution (sampling with replacement). Sampling without replacement is used to simulate the use of unique molecular identifiers that allow duplicate reads to be filtered.

### Performance assessment in simulations

We compared the performance of analysis methods for tiling CRISPR screens using several metrics. We computed the area under the precision-recall curve (prAUC) using scores provided by each method, specifically the functional sequence probability for RELICS, the −log10(FDR) for MAGeCK, the effect size (Beta) for CRISPR-SURF, and the Bayes Factor for BAGEL. Furthermore, we computed precision and recall using the set of regions labeled as significant by each method (BayesFactor > 5 for BAGEL, adjusted p-value < 0.05 for MAGeCK, positive significant regions output by CRISPR-SURF, and the functional sequences predicted by RELICS). Lastly, we also evaluated the fraction of all base pairs identified as significant that contained the simulated functional sequences.

### Running MAGeCK

MAGeCK (version 0.5.9.2) is designed to be run on sgRNAs that are grouped into functional units (i.e. genes). To run MAGeCK, we therefore grouped sgRNAs into non-overlapping genome windows containing 10 sgRNAs per window (note that the GeCKO v2 sgRNA library that MAGeCK is commonly applied to uses 6 sgRNAs per gene). To run MAGeCK we used the following command line:

~~~
mageck test -k MAGeCK_Input.txt -t 0,1 -c 2,3 -n mageckOut
~~~

where the MAGeCK_Input.txt file contains the observed counts for each sgRNA.

### Running BAGEL

BAGEL (downloaded from sourceforge on 12/12/2019) is designed to be run on sgRNAs that are grouped into functional units (i.e. genes). To run BAGEL we grouped sgRNAs using the same 10-sgRNA genomic windows that we used for MAGeCK. Since BAGEL takes log2 fold change as input, we calculated log2 fold change from the sgRNA counts. In addition, we specified a set of positive and negative sgRNAs to be used as training for BAGEL. For the simulations, the positive sgRNAs were taken from the simulated ‘positive control region’. As negative sgRNAs we used the first few 10-sgRNA windows which corresponded to the background and selected the same number of windows as was used as positive controls. To run BAGEL we used the following command:

~~~
python BAGEL.py -i BAGEL_Input.txt -o BAGEL_out.bf -e Input_posTrain.txt -n Input_negTrain.txt -c 1,2
~~~

### Running CRISPR-SURF

CRISPR-SURF (downloaded 02/13/2019 from GitHub) was run using the procedure specified in the supplemental materials of Hsu et al. 2018. To run CRISPR-SURF on the simulated datasets, we computed log2 fold change from the sgRNA counts and provided this as input to SURF_deconvolution. For the experimental datasets we filtered the sgRNAs and computed log fold change using the SURF_count command and then provided these log fold change estimates as input to SURF_deconvolution. The SURF_count command was run as follows:

~~~
docker run -v $Path_to_file/:$Path_to_file -w $Path_to_file pinellolab/crisprsurf SURF_count -f CRISPR_SURF_count.csv -nuclease cas9 -pert crispri
~~~

The SURF_deconvolution command was run as follows:

~~~
docker run -v $Path_to_file/:$Path_to_file -w $Path_to_file pinellolab/crisprsurf SURF_deconvolution -f CRISPR_SURF_Input.csv -pert crispri
~~~

### Running RELICS

To run RELICS on experimental datasets we used the default parameters and an sgRNA area of effect of 400bp (+/-200bp around each target site). The promoter regions of the genes of interest were used as positive control regions. The same positive control regions were used for CRISPR-SURF and BAGEL.

### CRISPRa screen data

We obtained data from published CRISPRa screens for *IL2RA* and *CD69* gene expression in Jurkat T cells (15). We downloaded the sgRNA counts from the paper supplement and converted them to the RELICS input format. The promoter regions were used as positive control regions for RELICS and CRISPR-SURF and were defined as the transcription start site (TSS) +/- 1kb. RELICS jointly analyzed all pools for the analysis. For MAGeCK and CRISPR-SURF, the input pool was used as control pool and the high-expression pool was used as the treatment pool. RELICS and CRISPR-SURF used the same positive control sgRNAs.

### CRISPRi screen data

We obtained data from published CRISPRi proliferation screens for *GATA1* and *MYC* in K562 cells (9). We downloaded the sgRNA counts from the paper supplement and converted them to the RELICS input format. For detecting *MYC* regulatory elements, all sgRNAs labelled as ‘*MYC* Tiling’ sgRNAs as well as ‘Protein Coding Gene Promoters’ sgRNAs were used. For detecting *GATA1* regulatory elements, all sgRNAs labelled as ‘*GATA1* Tiling’ sgRNAs as well as ‘Protein Coding Gene Promoters’’ sgRNAs were used. For MAGeCK and CRISPR-SURF, the pool at T0 (before) was used as the treatment pool and the pool at T14 (after) was used as the control pool in order to look for enrichment instead of depletion. RELICS and CRISPR-SURF used the same positive control sgRNAs.

### H3K27ac data

Jurkat H3K27ac data from Mansour et al.^34^ was downloaded from GEO (GSM1296384). The reads were aligned with BWA-MEM^35^ using default parameters and filtered for duplicates and low mapping quality (MAPQ < 30) using samtools^36^. The reads were then converted to reads per kilobase per million in 200bp bins using deepTools2^37^. ENCODE H3K27ac ChIP-seq data for the K562 cell line was downloaded in bedgraph format from the UCSC genome browser^38^.

### CRISPRa validation experiments

We used FlashFry (33) to design two sgRNAs per target site with high estimated efficiency and low estimated off-target effects (Supplementary table S1). The designed sgRNAs were cloned into pCRISPRiaV2 plasmids (Addgene #84832). Lentiviruses carrying the sgRNAs were generated and transduced into Jurkat cells expressing dCas9-VP64 (15), which were obtained from the Berkeley Cell Culture Facility. 7-10 days after transduction, cells from different treatments were stained with PE anti-human CD69 antibody (Biolegend #310906) and the expression of CD69 was measured by flow cytometry using a BD FACSCanto II system.

### CRISPRi validation experiments

K562 cells were transduced with dCas9-KRAB-GFP lentivirus (Addgene #71237) carrying different sgRNAs (Supplementary table S2). The percentage of GFP positive cells was recorded by FACS 3 days after transduction (D0) and again after an additional 6 (D6) and 14 (D14) days of culture. Two biological replicates (from separate cultures) were performed at each time point.

### RELICS model setup and organization

A description of the variables for the RELICS model is provided in Supplementary Table 3. RELICS divides the screened genome region into *M* small non-overlapping genome segments indexed as *m* = 1,2,…, *M*. In practice we have found a segment size of 100bp to work well, and we use this as the default. As input, RELICS takes a table of observed counts for *N* sgRNAs (indexed as *n* = 1,2,…, *N*) across *J* pools. We represent these counts as an *N* × *J* matrix, ***y***, and use ***y***_*n*_ to denote the vector of observed counts for sgRNA *n* across the *J* pools. Let *g*(*n*) be a function that maps sgRNAs to associated ‘overlapping’ genome segments. I.e. *g*(*n*) returns the set of genome segments that are overlapped by sgRNA *n*. RELICS does not use non-targeting sgRNAs.

### RELICS sgRNA count model

RELICS assumes there are *K* functional genome sequences, indexed as *k* = 1,2,…, *K* and that each functional sequence affects the sorting probabilities of overlapping sgRNAs. Each functional sequence has a length, *l_k_*, which is the number of genome segments that it spans. The maximum length of a functional sequence is set to *L*, so that *l_k_* ∈ {1,…, *L*}. By default, *L* = 10.

We describe the observed counts for an sgRNA across pools with a Dirichlet multinomial distribution and refer to the Dirichlet portion of the distribution as the sorting probability distribution. The sgRNAs that overlap a functional genome sequence have one sorting probability distribution, and the sgRNAs that do not overlap a functional genome sequence have another. Each Dirichlet distribution has *J* shape parameters, which define the dispersion and the expected proportion of counts in each pool. The vectors of shape parameters for the Dirichlet distributions are denoted ***α_1_*** and ***α_0_*** for sgRNAs that do and do not overlap a functional sequence. Then:

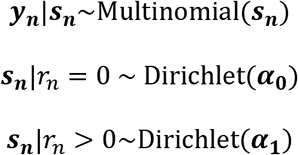

where *r_n_* is the total number of genome segments containing functional sequences that are overlapped by sgRNA *n*.

### RELICS functional sequence configurations

We define a configuration to be the positions and lengths of all of the functional sequences. We specify a configuration with a matrix, ***δ***, of dimension *K* × *M*, where an element, *δ*_***k,m***_, is 1 if genome segment *m* contains functional sequence *k*, and is 0 otherwise. We call a single row vector of the configuration matrix, ***δ_k_***, the placement of a functional sequence. In other words, a configuration is a collection of functional sequence placements.

We want to estimate the probability, *p_m_*, that a given genome segment, *m*, contains a functional sequence. To compute *p_m_* we could sum the posterior probabilities of all configurations that have a functional sequence in genome segment *m*. However, exact calculation of *p_m_* is intractable because the likelihoods of all possible configurations must be computed. For example, even in a simple case where there are *M* = 10,000 genome segments and *K* = 5 regulatory sequences of length *L* = 1, the number of possible ***δ*** configurations is 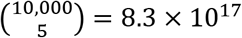.

To overcome this problem, we developed an approximate inference algorithm known as Iterative Bayesian Stepwise Selection (IBSS) (36). Our version of the IBSS algorithm includes extensions that are specific to RELICS including allowing for functional sequences of variable length and the use of non-normal count data with a Dirichlet multinomial error distribution. Our IBSS algorithm performs stepwise placement of a single functional sequence at a time, while accounting for the (uncertain) placements of all of the other functional sequences.

To implement the IBSS algorithm, and to account for uncertainty in functional sequence placements, we introduce a functional sequence probability matrix, ***π***. This is like the ***δ*** matrix, but rather than binary values, it contains probabilities. Specifically, each element is the probability that a genome segment contains a specific functional sequence: *π_k,m_* = Pr(*δ_k,m_* = 1).

To allow the positions of known functional sequences (positive controls) to be specified, we add an additional row to the functional sequence probability matrix, which we index as row 0 and denote ***π***_0_. The probability that a genome segment contains *any* functional sequence is then: 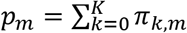.

The number of genome segments that are overlapped by sgRNA *n* and contain a functional sequence follows a Poisson binomial distribution. In other words, the Poisson binomial is used to calculate the probability that an sgRNA overlaps *r_n_* genome segments containing functional sequences:

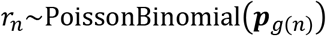

where ***p***_*g*_(*n*) is the vector of probabilities for all genome segments associated with sgRNA *n*.

### RELICS IBSS algorithm

The following is a description of the IBSS algorithm that is used to estimate the functional sequence probability matrix ***π***.

<u>Initialize</u>:

- Set *K* to number of functional sequences.
- Set known functional sequences (positive controls) to 1.0 in row ***π***_0_.
- Set all other elements of ***π*** to 0.0.

While not converged:

- Estimate sgRNA sorting hyperparameters (*α*_0_, *α*_1_) by maximum likelihood. Estimate configuration probabilities
- For *k* in 1… *K*:

○ Set elements of row ***π_k_*** to 0.
○ Compute ***p***, the probability each genome segment contains one of the other functional sequences
○ Set elements of ***π_k_*** by calculating posterior probability of every possible placement of functional sequence *k*, conditional on ***p***.

The algorithm is considered to be converged when the maximum absolute difference in the updated functional sequence probability matrix (max(abs(***π***′ – ***π***))) less than a defined threshold.

### RELICS posterior probabilities of functional sequence placements

To compute the posterior probability of each functional sequence placement, we first compute the likelihood of all possible placements of functional sequence *k*, taking into account the placements of all of the other functional sequences. We set the elements of ***π_k_*** to 0 and also set all elements of ***δ_k_*** to 0, except those corresponding to the functional sequence placement being considered, which are set to 1. We denote a specific placement as ***δ_k_***^<*m,l*>^, where *m* is the genome segment containing the start of the functional sequence, and *l* is the length of the functional sequence. That is, ***δ_k_***^<*m,l*>^, is a vector of 0s except for elements *m*..(*m* + *l* – 1), which are set to 1. We can compute the probability that a given genome segment contains a functional sequence as 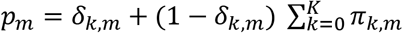. The likelihood of a specific functional sequence placement is then:

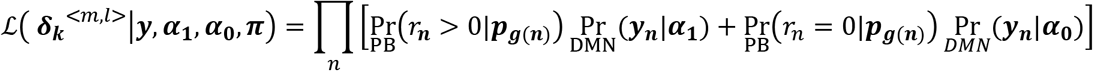

where the product is over all sgRNAs with observed counts, 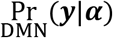 is the probability of the observed counts (computed using the Dirichlet multinomial distribution), and 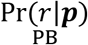 is the probability an sgRNA has *r* overlapping functional sequences (computed using the Poisson binomial distribution).

The posterior probability of a specific functional sequence placement is:

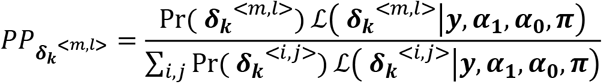

where the denominator is the sum over all possible placements of this functional sequence and Pr(***δ_k_***^<*m,l*>^) is the prior probability of a specific functional sequence placement.

We can compute the posterior probability that a genome segment contains functional sequence *k*, by summing the probabilities from all of the possible placements overlapping the segment. We use these posteriors to set the elements of ***π_k_***:

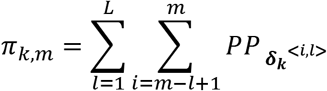

### RELICS prior probabilities of functional sequence placements

As prior probabilities for each functional sequence placement, we use a weighting that favors shorter functional sequences. Specifically, we use a geometric distribution, truncated at a maximum of *L*, to weight each possible length:

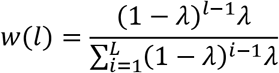

where *λ* is a constant between 0 and 1 that controls the weighting. We make the prior uniform for all placements with the same value of *l*.

### RELICS empirical estimation of hyperparameters

The RELICS model has hyperparameters, ***α*_0_** and ***α*_1_** which control the sorting probabilities and dispersion of sgRNA counts across pools. RELICS performs maximum likelihood estimation (MLE) of these parameters each iteration of the IBSS algorithm. This estimation is performed using the full dataset of sgRNA counts, and keeping the functional sequence probabilities fixed to their current estimates, 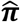:

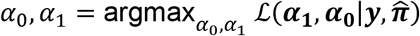

We perform MLE by numerical optimization using the L-BFGS-B algorithm (41).

### RELICS choice of the number of functional sequences (*K*)

RELICS estimates the number of functional sequences by first setting *K* to 1, and using the IBSS algorithm to estimate the functional sequence probability matrix, ***π***. *K* is then increased by one and the process is repeated. Each iteration we compute the pairwise Pearson correlations between all rows of ***π***. Initially the pairwise correlations are very low, however once the value of *K* is too high, the functional sequences placed by the algorithm begin to overlap causing positive correlations between pairs of functional sequence probabilities. In both simulations and real data, we have found it practical to set *K* to the largest value that yields a maximum pairwise correlation of less than 0.1. Figures 2, 3 and Supplementary Figures S5, S6 show how the maximum pairwise correlations increase dramatically once *K* exceeds the number of distinct functional sequences that can be identified from the data.

### RELICS Input Format

RELICS takes a text-based.csv file as input (for examples see https://github.com/patfiaux/RELICS). The file contains the sgRNA information as well as the observed sgRNA counts in different pools. We have made the public datasets we analyzed available in this format at (https://figshare.com/projects/RELICS_2_data/74376). Alternatively, RELICS can also take in two.csv files, where one file contains the sgRNA information and the other file contains the observed sgRNA counts in different pools.

### Availability of data and materials

RELICS is open source and available on GitHub (https://github.com/patfiaux/RELICS). All data were obtained from published papers or simulated as described above. The formatted data that we used for analyses can be downloaded from: https://figshare.com/projects/RELICS_2_data/74376.

## Acknowledgements

We thank Yarui Diao, Rongxin Fang, Ye Zheng, Zhi Liu, and members of the McVicker lab for helpful discussions about analysis methods for CRISPR regulatory screens; Arko Sen for guidance in obtaining and processing public datasets; Bing Ren, for his laboratory’s assistance with CRISPRi validation experiments; and Jessica Zhou and Arya Massarat for testing RELICS and CRSsim. This research was supported by NIH/NIAID grant 2R01AI107027-06; by NIH/NIDDK grant 1 R01 DK122607-01; by the National Cancer Institute funded Salk Institute Cancer Center (NIH/NCI CCSG: P30 014,195); by a gift from the Jacobs Foundation; by a fellowship from the H.A. and Mary K. Chapman Charitable Trust to P.C.F.; by a fellowship from the Jesse and Caryl Philips Foundation to P.C.F; by a Salk Alumni Fellowship to H.V.C; and by the Frederick B. Rentschler Developmental Chair to G.M.

## Author contributions

G.M. and P.C.F. conceived of the idea for RELICS. G.M. supervised the research. G.M. and P.C.F. wrote the manuscript, with input and edits from H.V.C. P.C.F. implemented RELICS and CRSsim, obtained the public data sets, and analyzed the data. H.V.C. participated in many helpful discussions about RELICS, CRSsim, and CRISPR screens. H.V.C. and A.R.C. performed CRISPRa validation experiments in Jurkat T cells. P.B.C. performed CRISPRi validation experiments in K562 cells.

## Competing interests

The authors declare no competing interests.

## Supplementary Figures

**Supplementary Figure 1.**
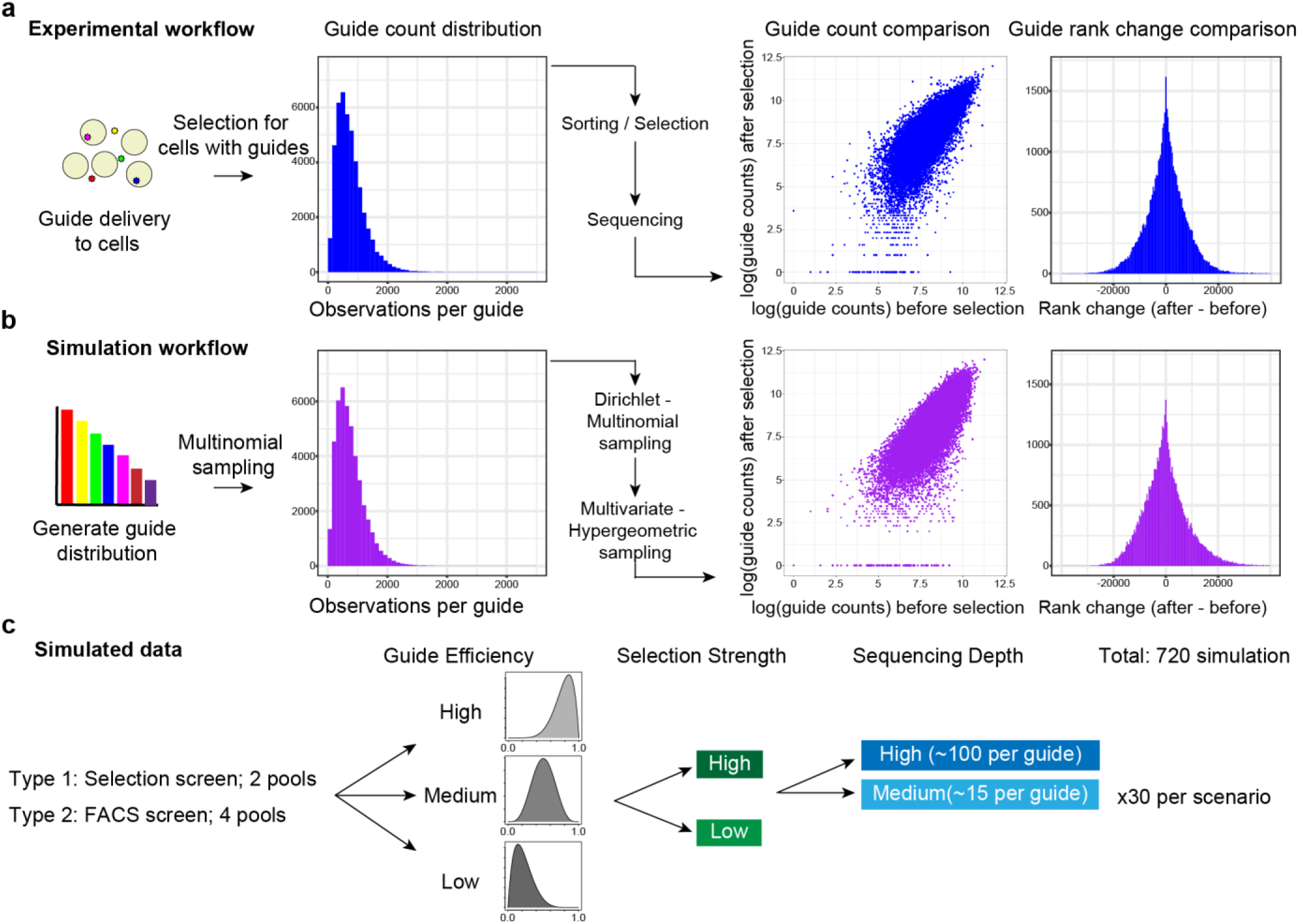
CRISPR screen simulation framework. **(a)** Experimental workflow. sgRNAs are introduced into cells and only cells which receive sgRNAs are retained. These cells are then either sorted based on gene expression, or placed under selective pressure (e.g. for survival or proliferation). sgRNA counts and guide ranks before and after selection are shown. **(b)** In our simulations, we generate an initial sgRNA count distribution and mimic the experimental steps using several different sampling procedures. The simulated sgRNA counts for before and after simulated selection are shown for comparison with the experimental data. **(c)** Combinations of parameters used for the simulations. We simulated two different types of screens with 3 different guide efficiency distributions, two different selection strengths, and sequencing depths. For each scenario, we simulated 30 data sets.

**Supplementary Figure 2.**
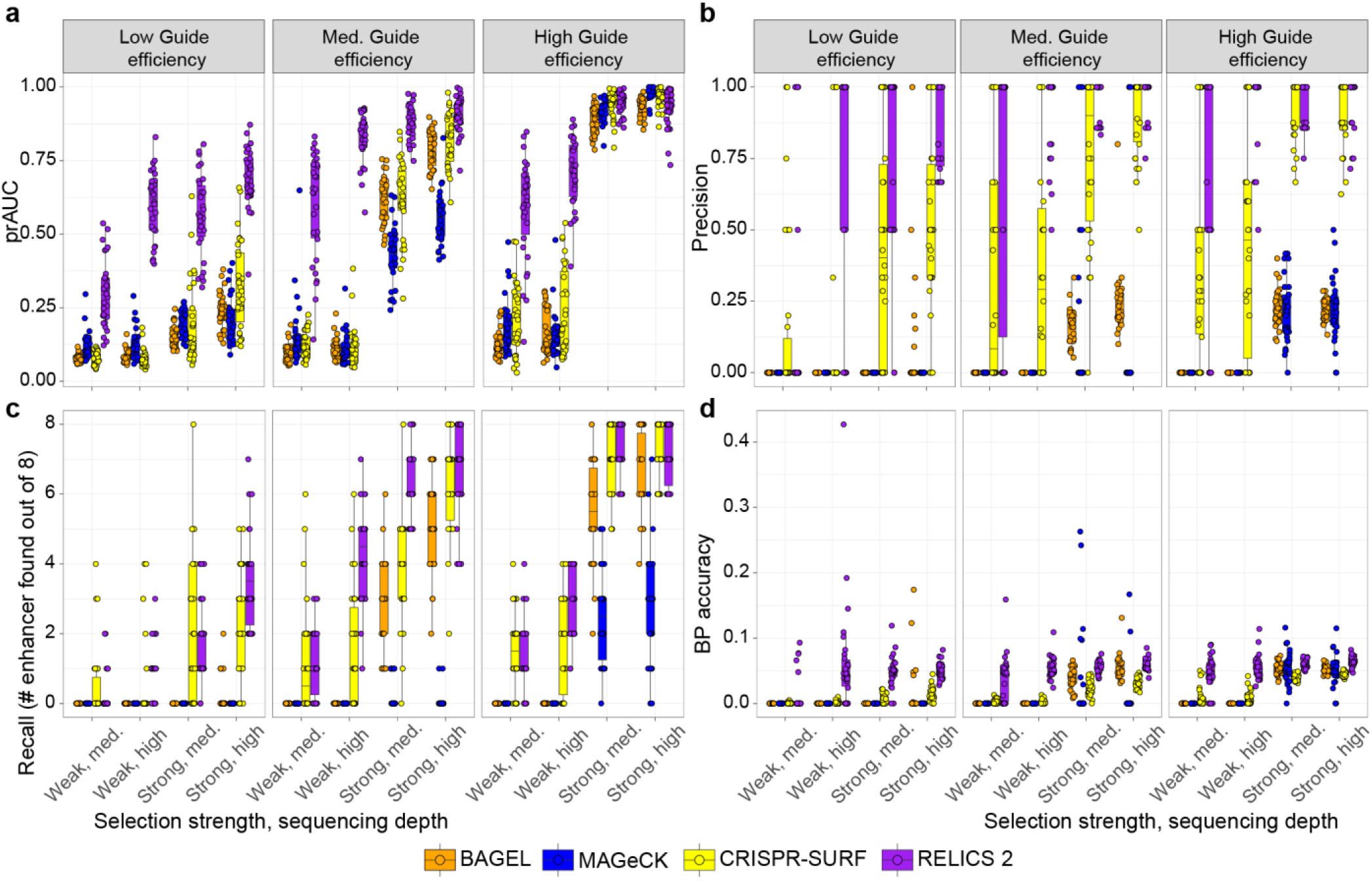
Performance evaluation of analysis methods on simulated data for a selection screen with 2 pools. **(a)** Boxplots of area under the precision recall-curve (prAUC) for different simulation parameters. **(b)** Boxplots of precision, the fraction of significant regions that contain a simulated enhancer. **(c)** Boxplots of recall, the number of simulated enhancers detected (out of 8) amongst the significant regions identified. **(d)** Boxplots of base pair (BP) accuracy, the fraction of base pairs in significant regions that overlap a simulated enhancer. The hinges of the boxplots correspond to the first and third quartiles, the center lines are the medians, and the whiskers extend to the furthest datapoints that are within 1.5x the interquartile range from the hinge.

**Supplementary Figure 3.**
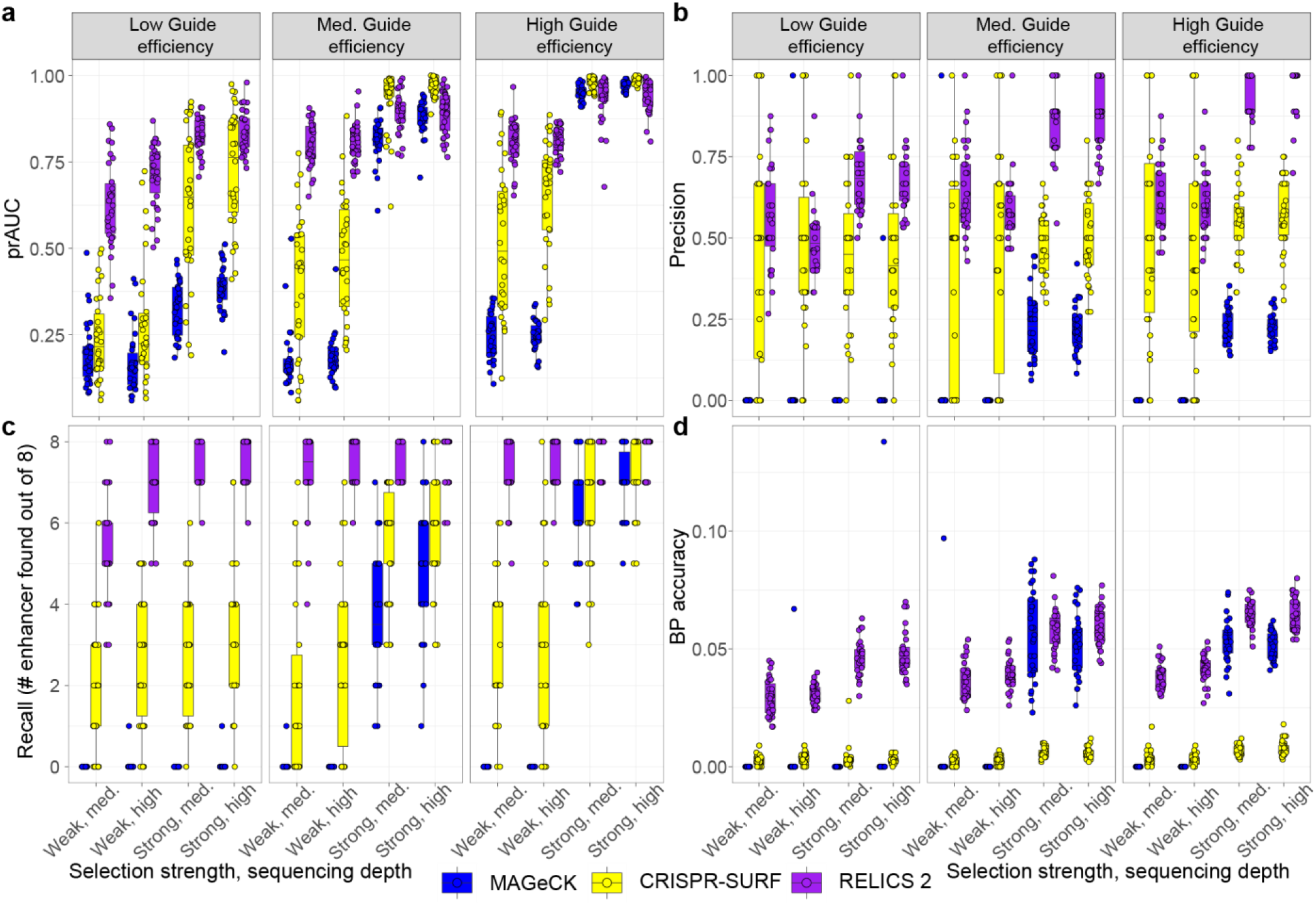
Performance evaluation of analysis methods on simulated data for a gene expression (FACS) screen with 4 pools. **(a)** Boxplots of area under the precision recall-curve (prAUC) for different simulation parameters. **(b)** Boxplots of precision, the fraction of significant regions that contain a simulated enhancer. **(c)** Boxplots of recall, the number of simulated enhancers detected (out of 8) amongst the significant regions identified. **(d)** Boxplots of base pair (BP) accuracy, the fraction of base pairs in significant regions that overlap a simulated enhancer. The hinges of the boxplots correspond to the first and third quartiles, the center lines are the medians, and the whiskers extend to the furthest datapoints that are within 1.5x the interquartile range from the hinge.

**Supplementary Figure 4.**
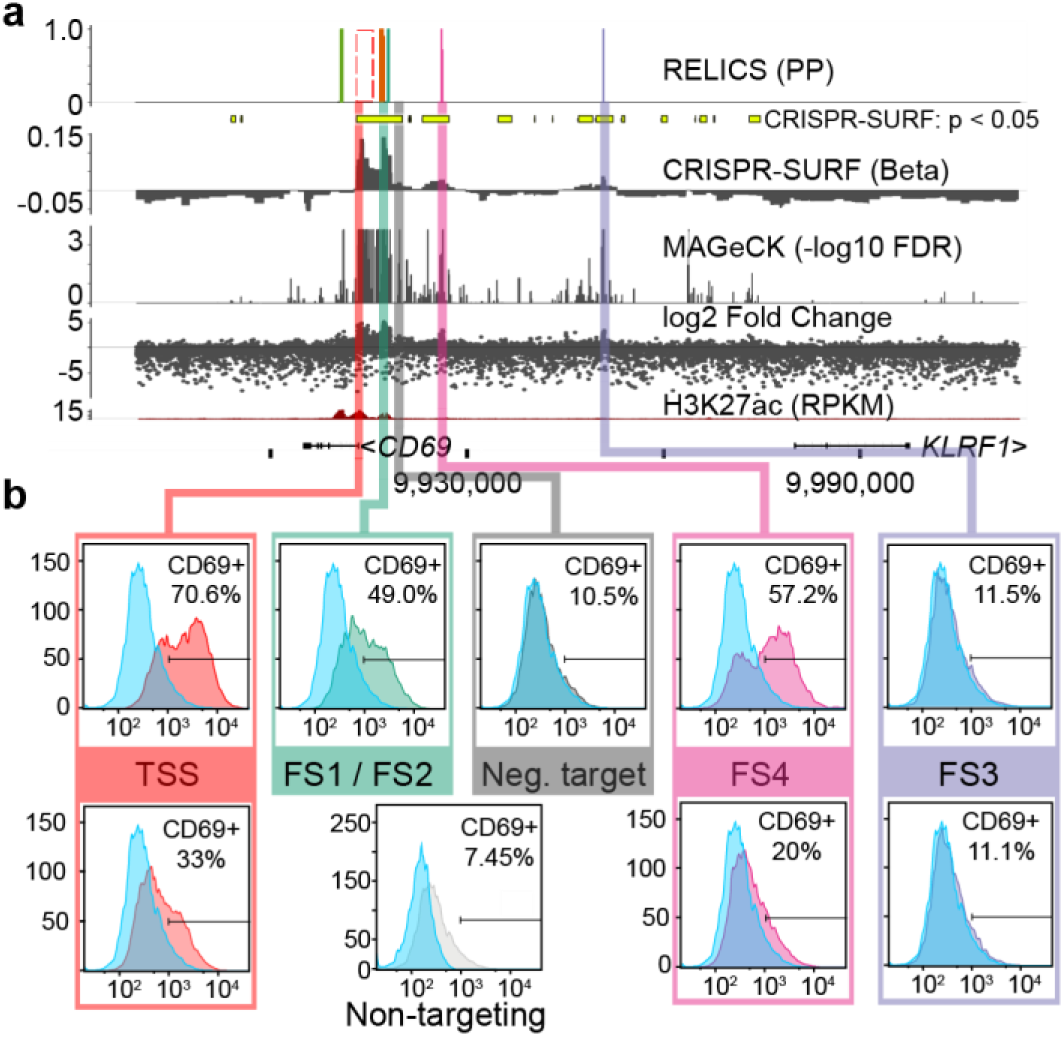
Experimental validation experiments for *CD69*. Lentiviral vectors encoding sgRNAs and dCas9:VP64 were co-transduced into Jurkat cells, and the expression of CD69 protein was quantified by flow cytometry. Non-targeting sgRNAs were used as a negative control. Targeting sgRNAs were chosen for their high predicted efficiency and in some cases are adjacent to the predicted FS rather than within the FS.

**Supplementary Figure 5.**
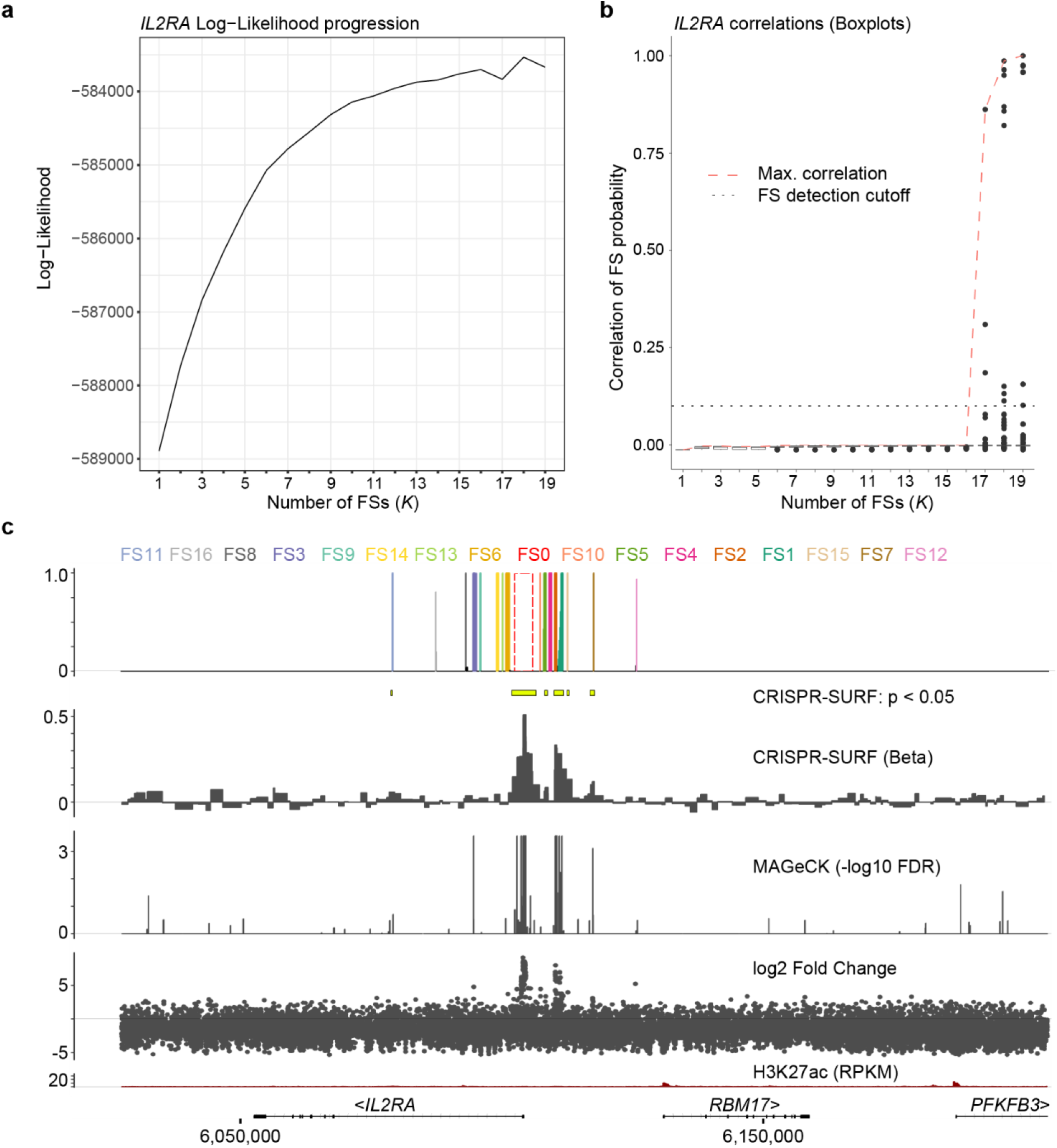
Analysis of a CRISPRa screen for *IL2RA* expression published by Simeonov et al. 2017. **(a)** Log-likelihood progression. With the addition of each functional sequence (FS), the model log-likelihood is recorded. **(b)** The pairwise correlations between functional sequence probabilities as a function of the number of FSs. The red dashed line indicates the highest pairwise correlation between all pairs of FSs. The hinges of the boxplots correspond to the first and third quartiles, the center lines are the medians, and the whiskers extend to the furthest datapoints that are within 1.5x the interquartile range from the hinge. **(c)** Output of RELICS and another analysis methods. Each FS predicted by RELICS is given a different color and the labels are arranged by genomic position.

**Supplementary Figure 6.**
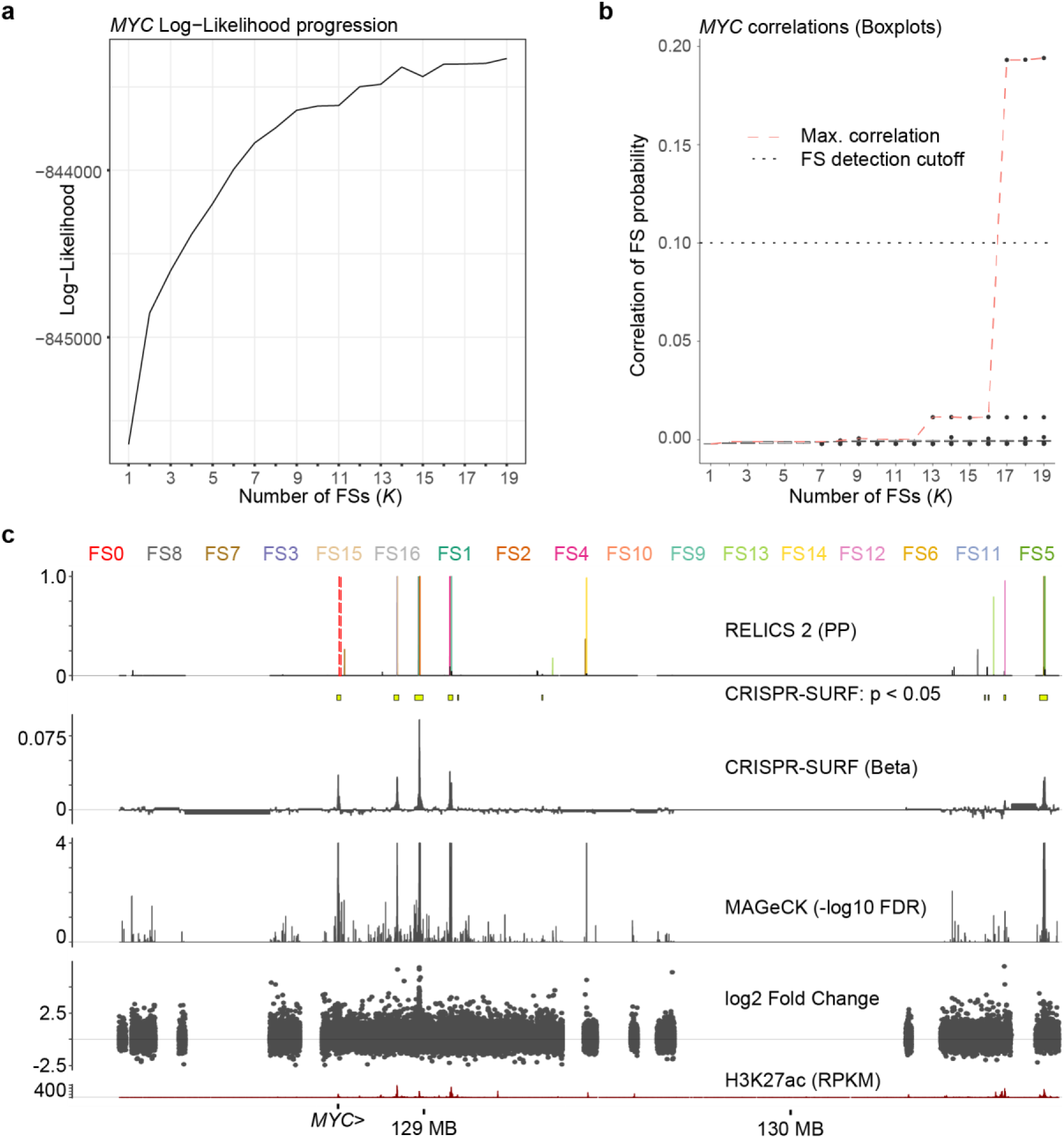
Analysis of a *MYC* CRISPRi cellular proliferation screen published by Fulco et al. 2016. **(a)** Log-likelihood progression. With the addition of each functional sequence (FS), the model log-likelihood is recorded. **(b)** The pairwise correlations between functional sequence probabilities as a function of the number of FSs. The red dashed line indicates the highest pairwise correlation between all pairs of FSs. The hinges of the boxplots correspond to the first and third quartiles, the center lines are the median, and the whiskers extend to the furthest datapoints that are within 1.5x the interquartile range from the hinges. **(c)** Output of RELICS and another analysis methods. Each FS predicted by RELICS is given a different color and the labels are arranged by genomic position.

**Supplementary Figure 7.**
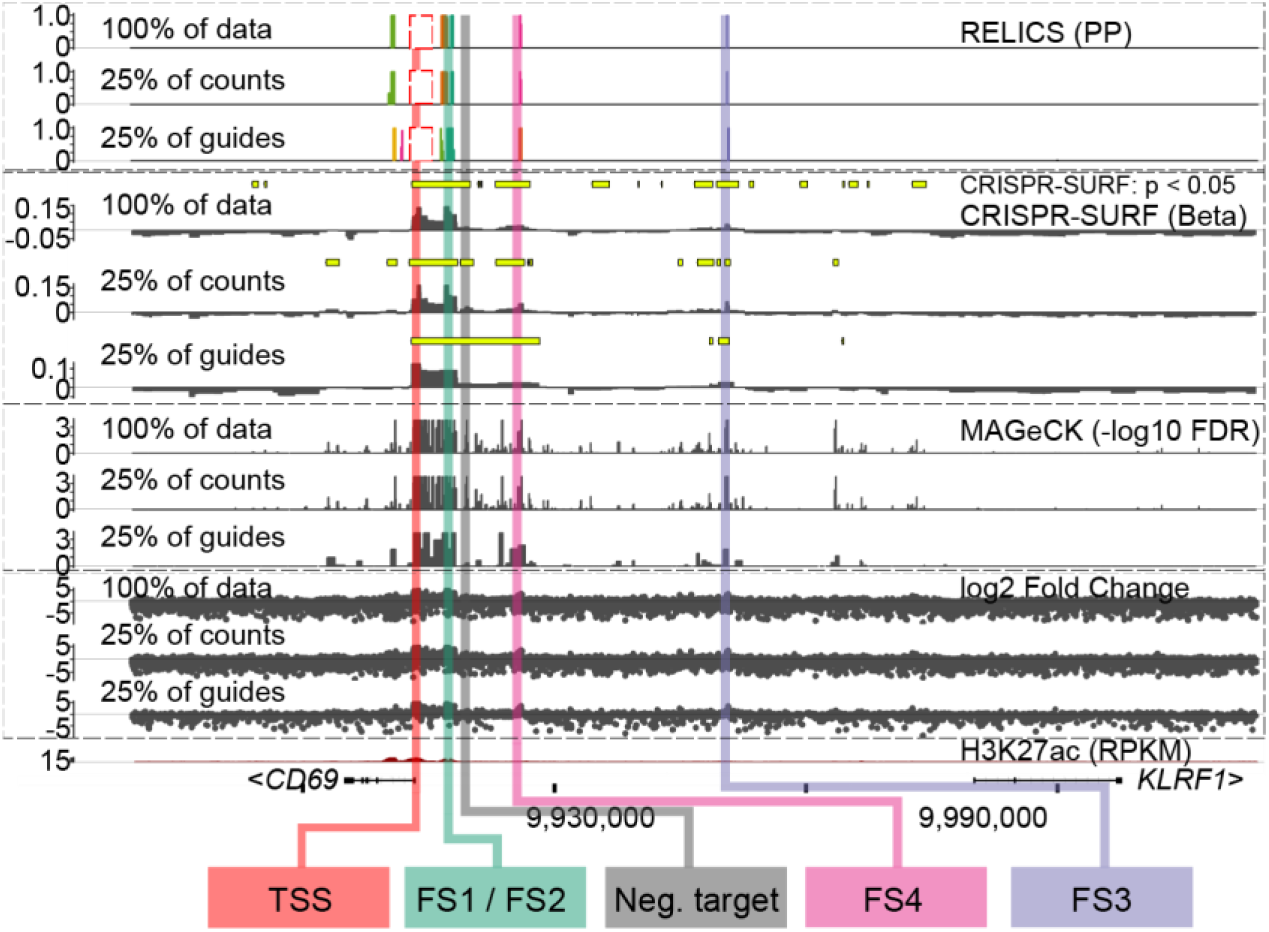
Downsampling of CD69 CRISPRa screen. sgRNAs counts were downsampled to 25% of all counts or sgRNA density was downsampled to 25%. Dashed blocks cluster results from different analysis methods together (RELICS, CRISPR-SURF, MAGeCK, log2 Fold Change).

## Supplementary Tables

**Supplementary Table 1.**
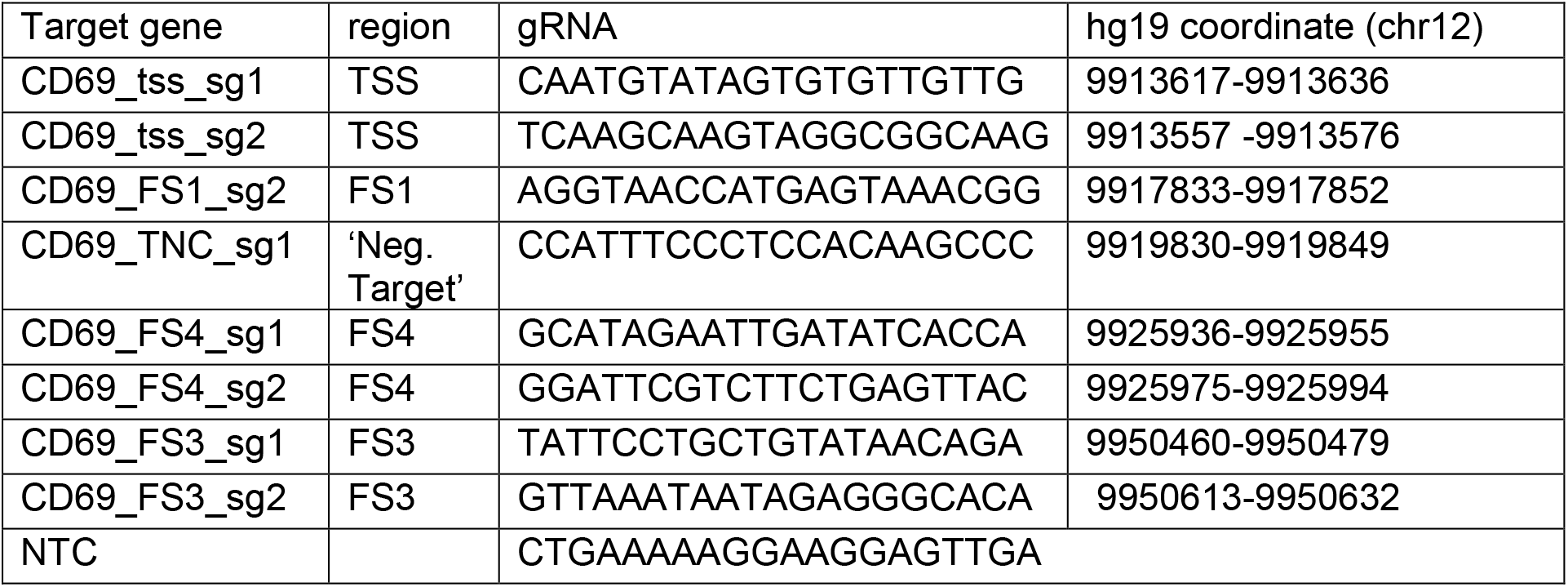
*CD69* CRISPRa validation sgRNAs. TSS sgRNAs target the transcription start site of *CD69*. FSs, and ‘Neg. Target’ sgRNAs correspond to labels in Fig. 4d. NTC is a non-targeting control sgRNA.

**Supplementary Table 2.**
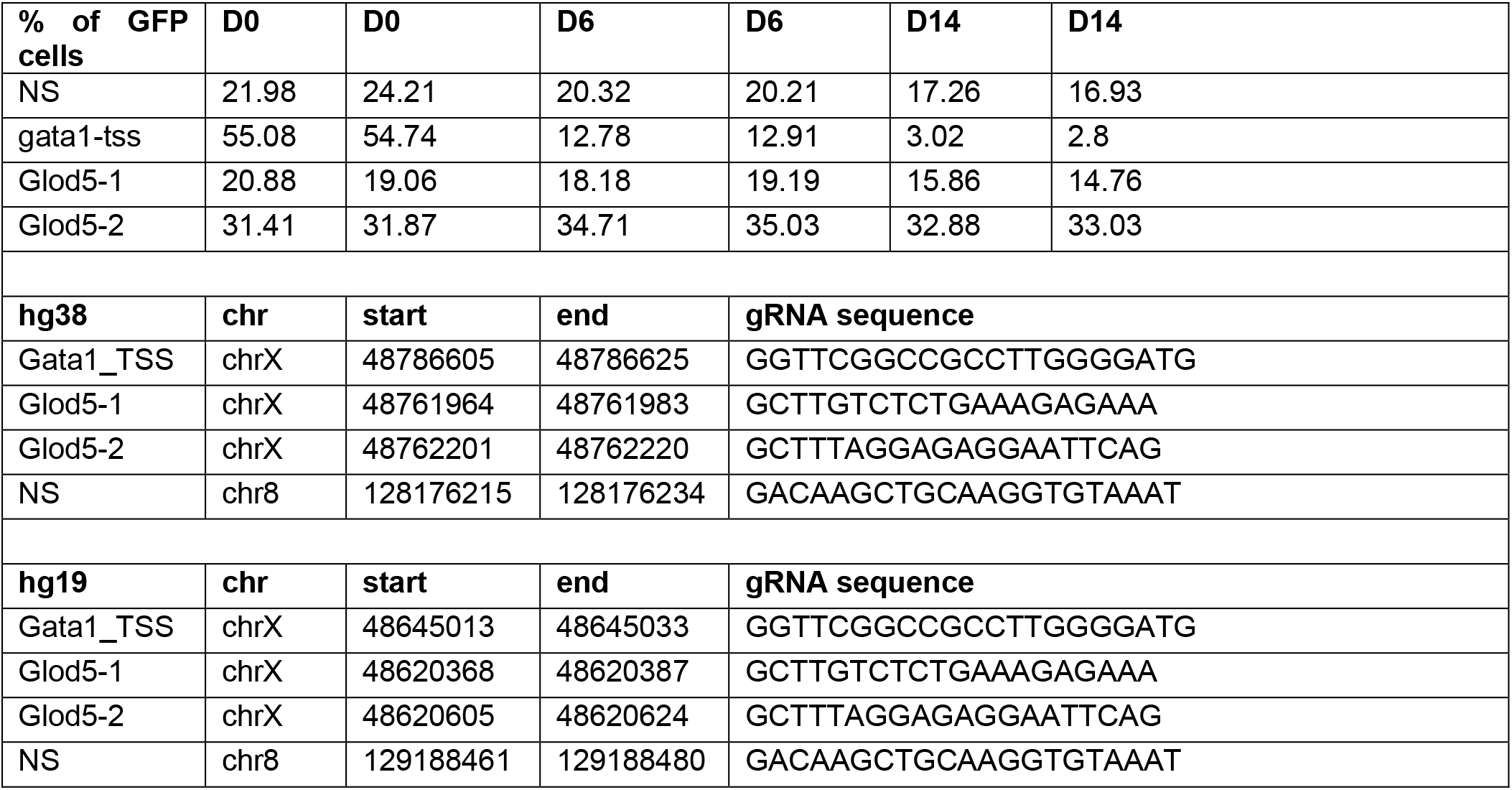
*GATA1* CRISPRi validation sgRNAs. Coordinates are given in both hg38 and hg19 genome assemblies. NS is a negative control sgRNA targeting a region on chromosome 8. GATA1_TSS sgRNA targets the transcription start site of *GATA1*. The Glod5-1 / Glod5-2 sgRNAs target the putative functional sequence near *GLOD5* identified by Fulco et al. 2016.

**Supplementary Table 3.** Description of variables in the RELICS model.

| Variable | Description |
| --- | --- |
| $J$ | Number of pools in screen |
| $K$ | Number of functional sequences |
| $L$ | Maximum length of functional sequence in genome segments |
| $M$ | Number of genome segments |
| $N$ | Number of sgRNAs |
| $y$ | Observed sgRNA counts (matrix of dimension $N \times J$) |
| $\delta$ | Functional sequence configuration (matrix of dimension $K \times M$). Each row, $\delta_k$ specifies the placement (length and position) of functional sequence $k$. A specific placement is denoted $\delta_k[m, l]$, where $m$ is the genome segment containing the start of the functional sequence and $l$ is the length of the functional sequence. |
| $\pi$ | Probability a genome segment contains a specific functional sequence (matrix of dimension $K \times M$) |
| $\alpha$ | Hyperparameters for sorting probability distribution (vector of length $J$) |
| $s$ | sgRNA sorting probabilities (vector of length $N$) |
| $r$ | Number of genome segments that an sgRNA overlaps that contain a functional sequence (vector of length $N$) |
| $p$ | Probability genome segments contains any functional sequence (vector of length $M$) |
| $l$ | Length of a functional sequence |
| $g(n)$ | Mapping of sgRNAs to genome segments |

## Notes

#### Summary of Updates

This version of the manuscript has been updated to include an improved implementation of RELICS which now includes the following: 1. RELICS is now a Bayesian hierarchical model and identifies one functional sequence at a time in a stepwise manner. 2. RELICS now is a semi-supervised method and updates its hyper-parameters with each additional functional sequence identified. 3. RELICS now considers the spatial organization and overlapping effects of multiple nearby sgRNA target sites and functional sequences. 4. RELICS provides interpretable probabilistic output for each functional sequence that can be used to delineate small genome regions that confidently contain each functional sequence. In addition to improved performance we have also included experimental validations of predicted functional sequences to highlight the accuracy of RELICS. All the above resulted in a completely new set of figures and more in-depth description of experimentally validated regions. In addition the author list has been updated to add two authors who contributed to the revised manuscript.

https://figshare.com/projects/RELICS_2_data/74376

https://github.com/patfiaux/RELICS

https://github.com/patfiaux/CRSsim

